# Phase-specific microstimulation in brain machine interface setting differentially modulates beta oscillations and affects behavior

**DOI:** 10.1101/622787

**Authors:** Oren Peles, Uri Werner-Reiss, Hagai Bergman, Zvi Israel, Eilon Vaadia

## Abstract

It is widely accepted that beta-band oscillations play a role in sensorimotor behavior. To further explore this role, we developed a novel hybrid platform to combine operant conditioning and phase-specific intracortical microstimulation (ICMS). We trained monkeys, implanted with 96 electrodes arrays in motor cortex, to volitionally enhance local field potential (LFP) beta-band (20-30Hz) activity at selected sites using a brain-machine interface (BMI). We demonstrate that beta oscillations of LFP and single-unit spiking activity increased dramatically with BMI training, and that pre-movement Beta-power was anti-correlated with task performance. We also show that phase-specific ICMS modulated the power and phase of oscillations, shifting local networks between oscillatory and non-oscillatory states. Furthermore, ICMS induced phase-dependent effects in animal reaction times and success rates. These findings contribute to unraveling of the functional role of cortical oscillations, and to future development of clinical tools for ameliorating abnormal neuronal activities in brain diseases.

## Introduction

Oscillatory activity, in a variety of frequency bands, is considered, by both theoretical and experimental neuroscientists, as a possible mechanism for the generation of assemblies of neurons that contribute to various functions from sensation to action and cognition (Gray et al., 1989; Hernandez-Gonzalez et al., 2017; Murthy and Fetz, 1996). Furthermore, these oscillations are also implicated in brain diseases (Brown et al., 2001; Raz et al., 2000; Uhlhaas and Singer, 2010), suggesting that they may reflect system abnormalities.

In this study, we focused on oscillatory activity in the beta frequency band of 20-30Hz (termed below as Beta) in motor cortex. Beta has been hypothesized to support synchronization over large distances (Kopell et al., 2000) and has been linked to different cognitive functions such as attention (Buschman and Miller, 2007) and working memory (Lundqvist et al., 2016). In motor cortex it has been demonstrated that Beta increases during hold periods, attenuates during movement initiation, and re-emerges thereafter (Baker et al., 1997; Pfurtscheller et al., 1996; Sanes and Donoghue, 1993; Tan et al., 2016). Engel and Fries suggested that Beta is related to preservation of the current sensorimotor or cognitive state (“status quo hypothesis”) (Engel and Fries, 2010). However, the functional relations between Beta, neuronal computation and behavior remain unclear.

There is also increasing evidence that patients suffering from neurological and psychiatric diseases exhibit abnormal oscillatory activity that may reflect impairment in neural dynamics. For instance, it has been established that Beta is greatly enhanced in Parkinson’s disease (Brown et al., 2001; Little and Brown, 2014; Raz et al., 2000), whereas a reduction in Beta and gamma oscillatory activity has been observed in schizophrenic patients during the execution of a wide range of cognitive tasks (Uhlhaas and Singer, 2010).

The lack of understanding of the role of Beta in health and disease has led to a growing interest in this frequency range and to increasing attempts to manipulate these oscillations and the neuronal activity that underlies them.

One approach is to exploit the plasticity of neuronal circuits – training the brain to modulate endogenously-generated oscillations by neural operant conditioning (biofeedback) (Fetz, 1969). Combining brain machine interfaces (BMI) with neural conditioning led to experiments in which subjects were trained to modify their brain activity in order to initiate actions and receive rewards. In a previous study conducted in our lab, it has been demonstrated that the power of local field potential (LFP) in the gamma band (30-43Hz) can be enhanced using this paradigm (Engelhard et al., 2013). Recently, Beta has also been the target of similar operant conditioning schemes, which demonstrated positive correlation between the volitionally induced Beta-power and the reaction time (RT) (Khanna and Carmena, 2017; Peles et al., 2016).

An additional classic intervention tool, is the intracortical microstimulation (ICMS) (Stoney et al., 1968), which has been used extensively to modulate neuronal activity. In motor cortex, ICMS can evoke movements (Graziano et al., 2002), affect functional connectivity (Jackson et al., 2006), and disrupt or delay voluntary movements (Churchland and Shenoy, 2007; Griffin et al., 2011). In most studies, the ICMS has been used in open-loop conditions that did not depend on the ongoing activity.

Here, we use a closed-loop system, by detecting oscillations and examining the effects of ICMS on the detected pattern. The system is based on well-established evidence that during oscillations, some neuronal ensembles increase or decrease their firing rate at specific phases of the oscillation (Martin and Schroder, 2016; Murthy and Fetz, 1992). We assumed that stimulating at one of these preferred or non-preferred phases might change the balance between depolarization and hyperpolarization of neurons near the stimulation site. Careful choice of a specific stimulation phase could therefore induce opposite perturbations, enhancing an existing oscillation or alternatively, suppressing the oscillation. This notion is supported by recent computational (Holt et al., 2016), experimental (optogenetic stimulation (Siegle and Wilson, 2014) and transcranial magnetic stimulation (Guerra et al., 2016; Schilberg et al., 2018)) and clinical studies (deep brain stimulation (DBS) locked to the patient’s tremor (Cagnan et al., 2017)). A recent study conducted on Parkinson’s disease patients used open-loop deep brain stimulation, and applied offline analysis to demonstrate that the suppression of Beta was dependent on the phase relations between the electrical pulse and the neuronal oscillations (Holt et al., 2019). This promising result emphasizes the need for advanced methodologies for modulation of spatiotemporal patterns of brain activity. In particular, real-time detection of targeted neuronal patterns and closed-loop stimulation is one of the main challenges of translational, state-of-the-art studies (Holt et al., 2019; Holt et al., 2016; Moll and Engel, 2017; Zanos et al., 2018). In addition, advances in these methodologies will facilitate deeper understanding of the relation between experimental modulation, neuronal state, and behavior.

Here, we employed a novel technique, which combined operant conditioning and ICMS as synergistic approaches for manipulating cortical oscillations. We used a real-time BMI platform to induce volitional increase of Beta oscillatory activity, by reinforcing emergence of these patterns. Then while Beta was increased, we applied phase-specific ICMS at local sites in the primary motor cortex to explore the effects on the local network dynamics and the animal’s behavior.

## Results

Details of the experimental design are described in the Methods section. Briefly, two macaque monkeys were chronically implanted with arrays of 96 recording and stimulating microelectrodes (Blackrock Microsystems, Salt Lake City, UT, USA) in the hand area of motor cortex (see Figure S1). Data for this report include 2790 well-isolated single-units recorded in 80 daily sessions and LFP from 149 sessions. During each session, monkeys performed a visuomotor task while they volitionally controlled the Beta by means of neural operant conditioning.

Figure 1A shows the trial sequence. The monkey initiated a trial by moving its hand in response to a visual cue (orange circle, stage 0). LFP signals recorded by 2-3 *conditioning electrodes* in the array were used as a real-time input to the BMI algorithm. A visual signal in the form of a green ring (stage 1) was presented to the monkey, with its radius proportional to the Beta*-*power (20-30Hz). The Beta had to reach a minimal level *(threshold-reach*) before proceeding to the next stage of the trial, at which a visual cue (yellow or red circle) was presented to the monkey (stage 2).

**Figure 1.**
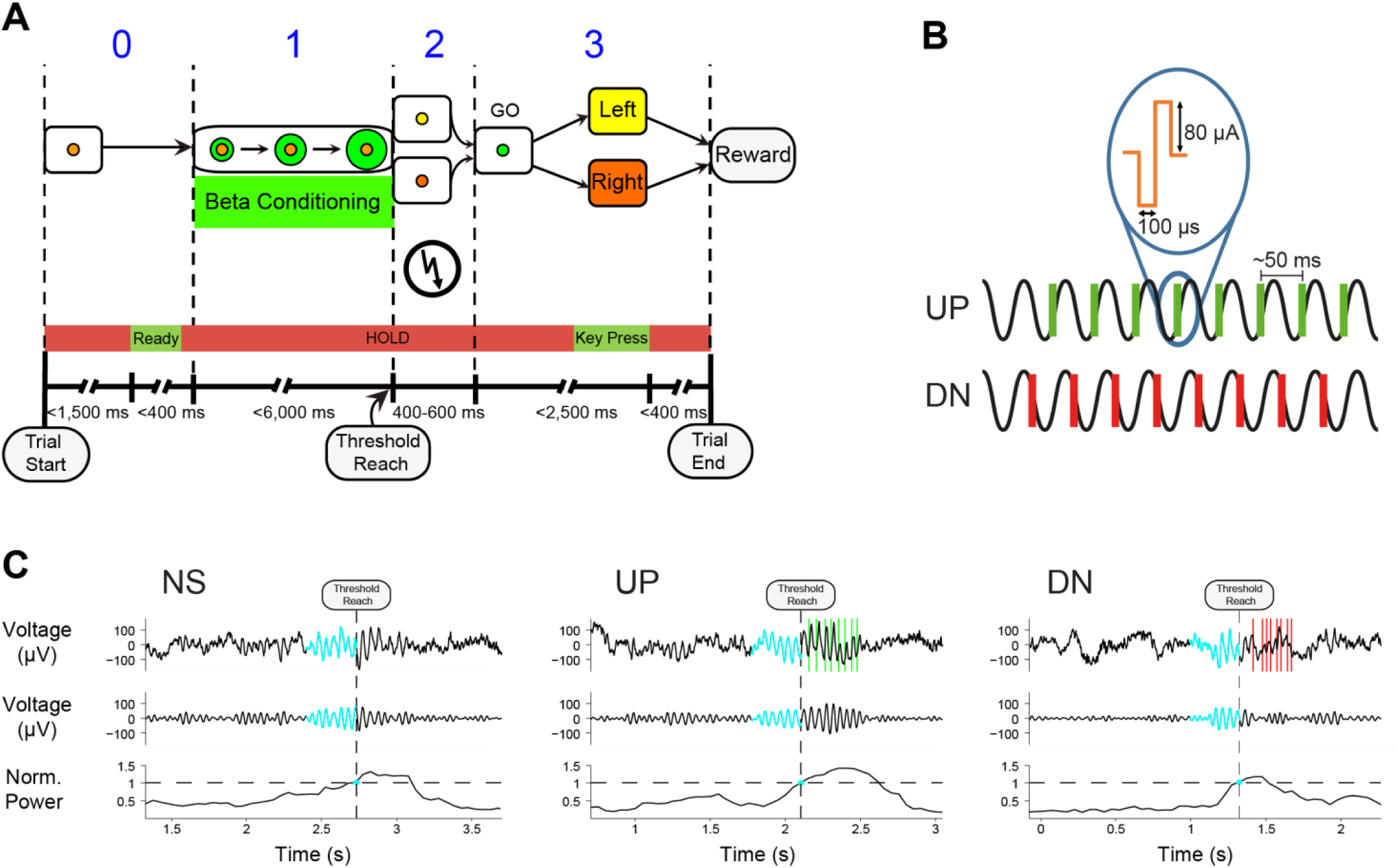
Trial scheme: behavior, recording and stimulation. (A) Trial scheme. Stage 0: The monkey responds to a cue (orange) to express alertness (“ready”). Stage 1: Beta enhancement conditioning. Stage 2: Visual cue (red or yellow), with or without ICMS (lightning symbol). Stage 3: Go-signal followed by key press. (B) ICMS properties with UP/DN stimulation illustration. (C) typical examples of NS, UP, and DN trials. Top panel: raw LFP signal averaged over the conditioning electrodes. In cyan, the epoch during which the Beta-power threshold was crossed. Vertical green and red lines mark the UP and DN ICMS-trains respectively. Middle panel: filtered LFP signal (20-30Hz). Bottom panel: Beta-power normalized by the power at *threshold-reach*. Threshold crossing point is at the intersection of the dashed black lines.

To explore how perturbations of a local network in an oscillatory state modulate the network dynamics, we applied ICMS to a single electrode located in the vicinity of the conditioned site of the array. ICMS was applied in randomly selected trials after *threshold-reach* (stage 2) and consisted of a train of 5 to 8 balanced bi-phasic pulses, termed as an ICMS-train (Figure 1B). An ICMS-train was precisely timed at the rising or falling phase of the LFP oscillation (hereafter referred to as “UP” and “DN” trials, respectively). Trials with no stimulation are referred to as “NS” trials. In all trials, to obtain a reward, oscillations had to be maintained until the Go-signal (circle turned green) and the monkey had to correctly respond to the visual cue by pressing the right or left key (stage 3).

Figure 1C shows single-trial examples of the LFP input to the BMI in NS, UP and DN trials. The top panel shows the raw LFP with the segment that reached the threshold marked in cyan. Vertical green and red lines mark the ICMS-train for UP and DN trials respectively. The middle panel shows the same trace filtered between 20-30Hz. The bottom panel shows the Beta-power with the desired threshold and the crossing point (cyan dot).

### Enhancing Beta by neural operant conditioning

The experiments began with a learning period (first 26 and 19 recording sessions for monkeys C and A respectively) during which the monkeys acquired and gradually improved volitional control of Beta oscillations. Figure 2 presents data from the learning period to illustrate this improvement of monkey C (left column) and monkey A (right column). All other analyses in this report are based on data from the rest of the sessions, after the monkeys reached stable volitional control of Beta.

**Figure 2.**
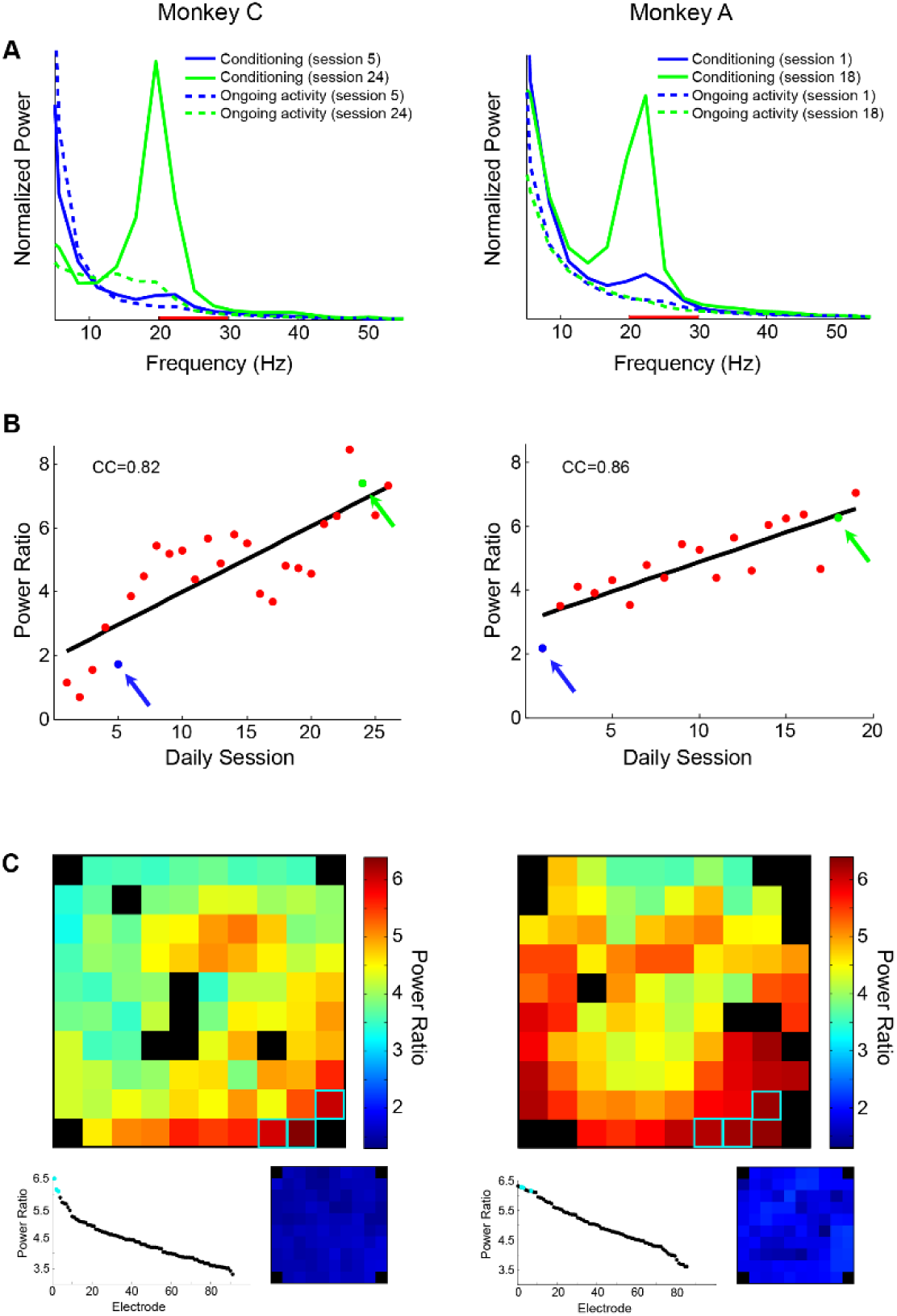
Enhancing Beta by neural operant conditioning. (A) Mean normalized power spectrum for monkey C (left) and A (right) at single recording days from early (blue) and late (green) sessions of the learning-period (first 26 and 19 sessions for monkeys C and A, respectively). The LFP power at the conditioning electrodes was computed over a 350ms epoch, for on-going activity (dashed line, starting 1s before trial-start), and for the conditioning (solid line, at threshold-reach). Red bar marks the conditioned frequency band. (B) Red dots denote the ratios between mean Beta-power during the conditioning and during on-going activity, across the learning period. Blue and green arrows and dots mark the early and late sessions (shown in (A)), respectively. Correlation coefficient (CC) between the variables was 0.82 for monkey C and 0.86 for monkey A. (C) Color-coded matrix showing Beta-power ratios (calculated as in (B)) for each of the 96 electrodes across the array at the end of the learning period. Conditioning electrodes are marked by cyan frames. Black squares represent noisy or non-functional electrodes. Bottom left: A graph showing sorted Beta-power ratios for all electrodes, with the conditioning electrodes marked in cyan. Bottom right: Beta-power ratios for the beginning of the learning period (small blue matrix).

Figure 2A shows the normalized power spectrum at the beginning (blue) and end (green) of the learning period, for on-going activity (dashed line, starting 1 second before *trial start)* and for the conditioning period (solid line, before *threshold-reach*). Figure 2B shows the ratio between the Beta*-*power during the conditioning period and during the on-going epoch, across daily sessions. The figure clearly demonstrates that the two monkeys learned to increase the Beta-power during the neural operant conditioning.

The color scaled matrices in Figure 2C display the power-ratios across the entire array at the beginning (small blue matrix) and at the end (large matrix) of the learning period. The figure shows that while there was little or no Beta on all electrodes at the beginning of learning, it increased dramatically by the end of learning around the conditioning electrodes (marked in cyan frames), and to some extent on all electrodes. This is also shown in the graph at the bottom left of the large matrix, where all electrodes are sorted by their post-learning power-ratios (conditioning electrodes marked in cyan).

### Beta oscillations of single-units exhibit phase locking to the LFP

During LFP Beta-conditioning epoch of each trial (Figure 1A, stage 1), we observed increased Beta in spiking activity, phase locked to the LFP Beta (Figure 3). Figure 3A presents examples of typical locking of spikes to the LFP oscillations. The top panel shows LFP from three electrodes averaged over one third of the trials with the highest Beta-power, and triggered on the last peak of the oscillation before *threshold-reach* (time=0). The three raster displays and peri-stimulus time histograms (PSTHs) of the lower panels show spiking activity of three neurons, recorded by the same electrodes and locked to the same time.

**Figure 3.**
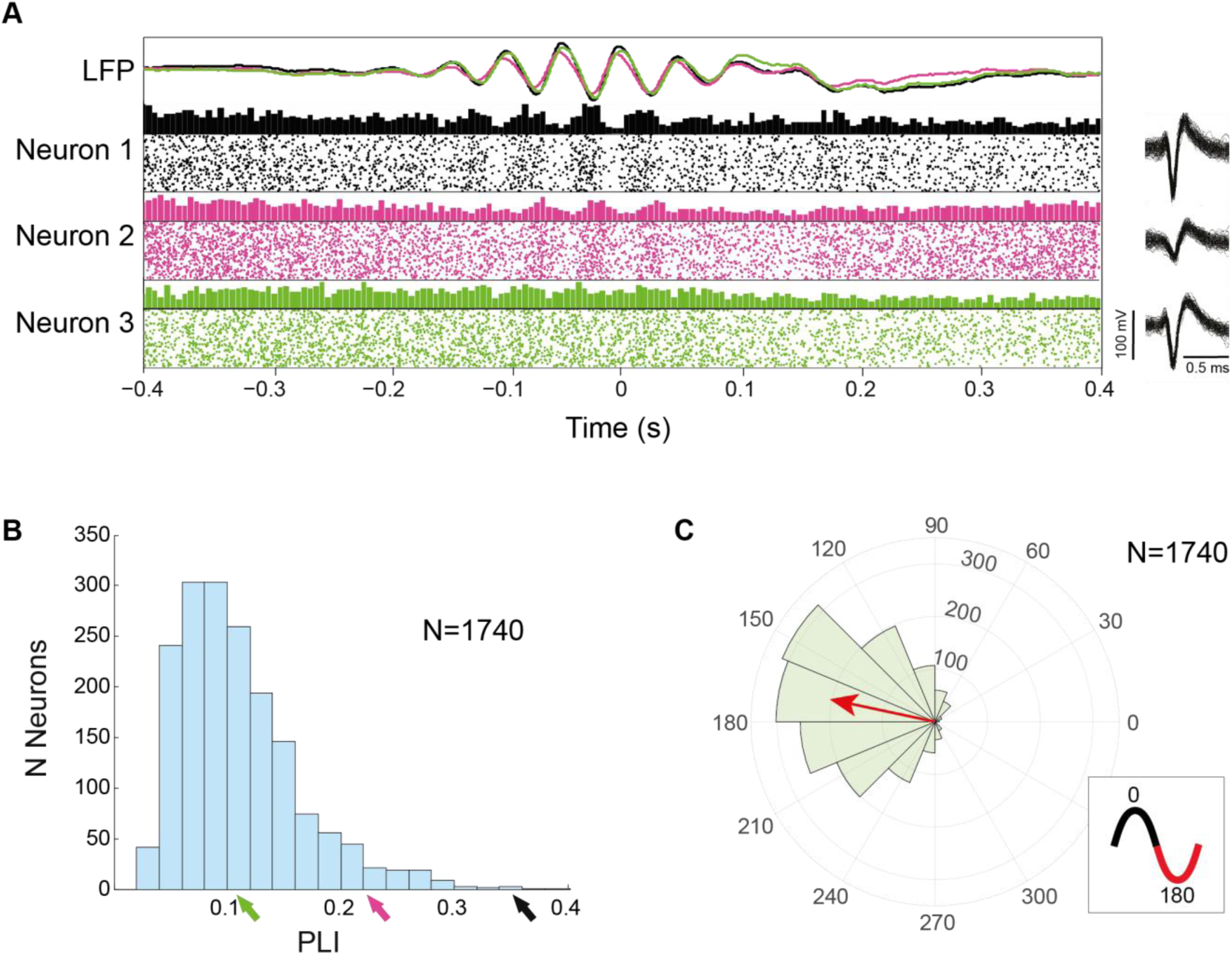
Beta Oscillations of single-units exhibit phase locking to the LFP. (A) Phase locking examples of three units. The top panel shows the mean LFP of three electrodes during conditioning, and the bottom panels, examples of three units recorded by the same electrodes. The raster displays and PSTHs for the three units depict varied degrees of phase locking. Time zero is the last positive LFP peak before *threshold-reach*. Right: 100 overlaid shapes of randomly selected spikes, of each unit. (B) PLI distribution of 1740 neurons having significant PLIs (p<0.01, bootstrapping, see Methods). Arrows mark the three units presented in (A). (C) Preferred-phase distribution for the significant-PLI neurons. Almost 90% of the neurons preferred the half cycle around the negative peak of the oscillation (marked in red in the inset cycle plot). Red arrow denotes the mean preferred phase. Data in (B) and (C) are based on 80 sessions.

To assess the LFP and spike synchrony, we calculated for each neuron a Phase-Locking Index, termed PLI, (ranging from 0 for no locking to 1 for perfect locking) and the preferred phase (see Methods). More than half of the sampled neurons (1740 of 2790) had statistically significant PLIs (p<0.01, bootstrapping, see Methods). For many of these neurons, the PLIs, while significant, were quite small (51% had PLI < 0.1) reflecting the variable nature of the relation between LFP and spikes firing. Figure 3B shows the distribution of significant PLI values. The three arrows on the x-axis mark the PLIs (0.1, 0.22 and 0.35) of the neurons in 3A.

Next, we explored the distribution of the preferred phases of neurons with significant PLI (Figure 3C). Note that only 10.2% of the neurons had their preferred phase during the half cycle around the positive peak of the oscillation, while almost 90% lie around the trough (90° to 270°, marked in red in the inset cycle plot of 3C). The figure also shows that the majority of neurons (60%) tend to synchronize their firing to the falling phase (0° to 180°) of the LFP Beta with the mean preferred phase being 168°.

To sum up, in spite of the low PLI value of each neuron, their tendency to fire in a limited phase-regime (mainly around the LFP trough) results in synchronous state of the network, as demonstrated in Figure 6A below (especially the epoch before time zero).

### LFP Beta-power before movement is anti-correlated with task performance

In order to explore how the monkey’s behavior is influenced by the volitionally enhanced Beta, we examined the monkey’s reaction time (RT, from Go-signal to movement initiation), movement time (MT, from movement initiation to key press), and success rate (correct key presses).

RT and MT were computed based on the signal of an accelerometer attached to the monkey’s hand and the key press time. Figure 4A shows two trial examples, depicting the accelerometer signal (blue) and the normalized Beta-power (red). Figure 4B shows that the RT+MT epoch and success rate were correlated with Beta-power, with a significant increase of the period from Go-signal to key press (RT+MT), and decrease of success-rate as Beta-power rises. The power was measured in a segment of 350ms around the Go-signal. These results, showing that Beta interferes with motor execution, are consistent with the results of previous studies (Joundi et al., 2012; Khanna and Carmena, 2017; Pogosyan et al., 2009) and the “status quo hypothesis” (Engel and Fries, 2010).

**Figure 4.**
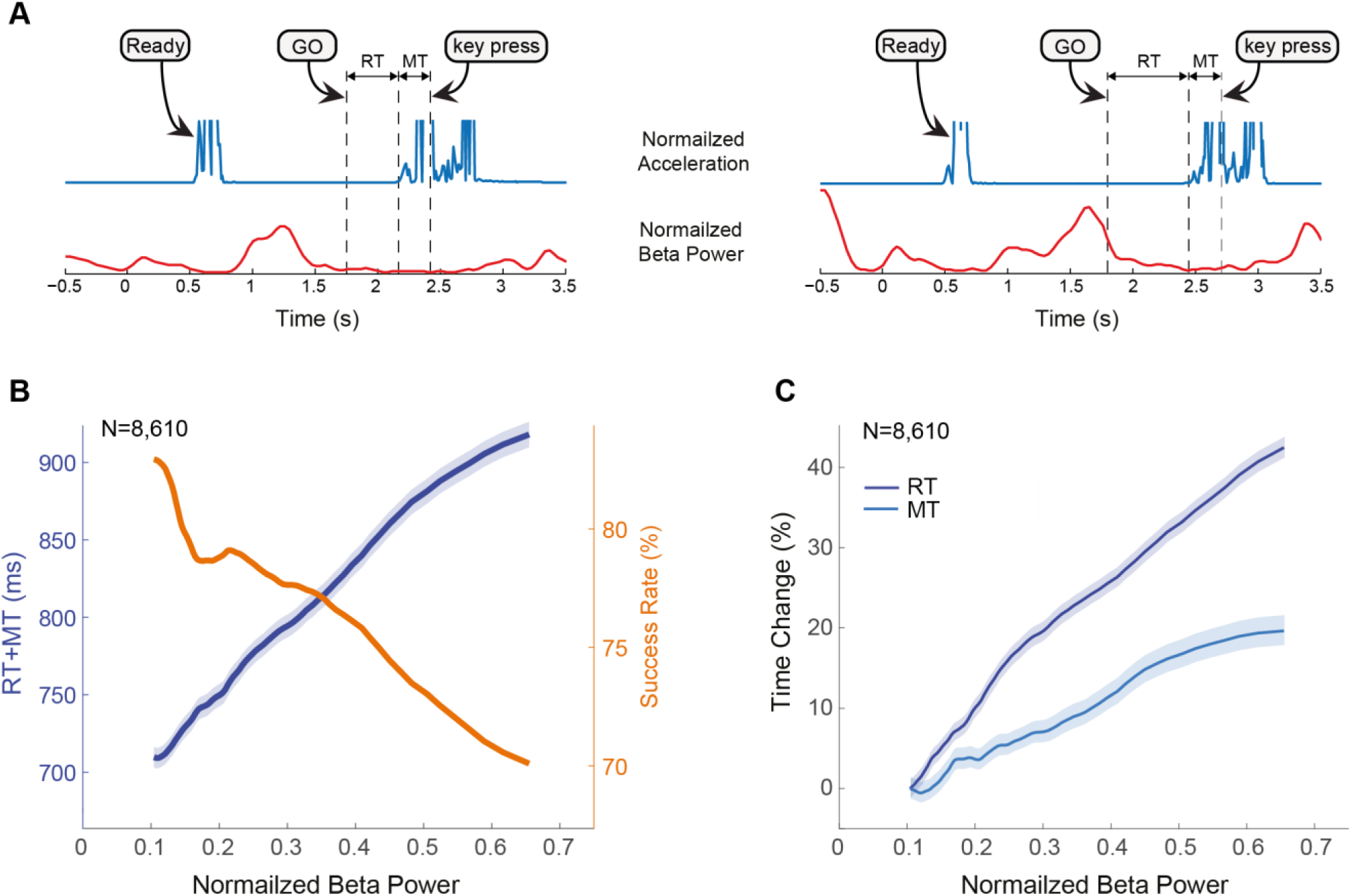
LFP Beta-power before movement is anti-correlated with task performance. (A) Examples of two single trials with shorter (~400ms) and longer (~650ms) RTs, showing the accelerometer signal (blue graph, used to compute the RT) and the normalized Beta-power (red graph). Dashed vertical lines mark (from left to right) the Go-signal, the movement onset and the key press. “Ready” marks the monkey’s initial movement to express alertness. Accelerometer signal is magnified and truncated to highlight movement initiation after Go-signal. Time zero is the beginning of the trial. (B) Success rate (orange) and RT+MT (blue) vs. Beta-power, normalized by the power at *threshold-reach*. (C) Change in RT (dark-blue) and MT (light-blue) vs. the normalized Beta-power. The change is relative to the RT and MT at the lowest Beta point (at ~0.1). Mean RT and MT are 507ms and 276ms respectively. The graphs show moving average with SEM. Significance was tested between the first (lowest Beta) and last (highest Beta) points of the graphs (p<0.001, Fisher’s test for success rate and Wilcoxon signed-rank test for RT, MT and RT+MT). Data are based on NS (no stimulation) trials from 30 sessions.

It has been debatable whether the Beta-related slower response is due to a longer reaction time (RT) or a slower movement. Pogosyan et al. reported that when Beta was induced by transcranial alternating current stimulation (tACS), the movement time increased, but RT was unaffected. Khanna et al., on the other hand, showed that following volitionally induced Beta, the RT decreased, but movement velocity-parameters were not correlated with the Beta-power. Here we found that both RT and MT were longer in high Beta-power trials (Figure 4C), but partial-correlation analysis reveals that the statistically significant partial correlation is that of Beta-power and RT (Beta-RT: CC=0.247, p<0.0001, Beta-MT: CC=0.027, p=0.025) in accordance with Khanna et al. We therefore use only RT hereafter to evaluate the relations between Beta and motor behavior.

### Differential effect of stimulation phase on LFP Beta

Once the monkeys learned to volitionally enhance Beta-power, we explored how perturbations of a local network in an oscillatory state modulate the network dynamics. To do this, we applied ICMS to a single electrode located in the vicinity of the conditioned site of the array. Figures 5A-F describe the effect of ICMS on the LFP at the conditioning electrodes. The plots show that stimulating at the rising phase of the oscillation (UP, green) increased the Beta while stimulating at the falling phase (DN, red) decreased or did not affect the oscillations as compared to the oscillatory activity in trials with no stimulation (NS, blue). The difference can be seen at the single-trial level for UP (Figure 5A) and DN (Figure 5B) trials, as well as at the average activity (Figure 5C). Note that the ICMS pulses in UP and DN trials fall on opposite phases of the LFP, indicating accurate phase detection. The plots in Figure 5D display histograms of the Beta-power ratio between the Beta during stimulation and the Beta before *threshold-reach*. While the oscillations tend to decrease in NS trials (power ratio<1 in 77.4% of the trials, median ratio of 0.52), they significantly increase in UP trials (power ratio>1 in 70.6%, median ratio of 1.53). In contrast, during DN stimuli the Beta-power decreased even below the NS trials (power ratio<1 in 87.6%, median ratio of 0.34).

**Figure 5.**
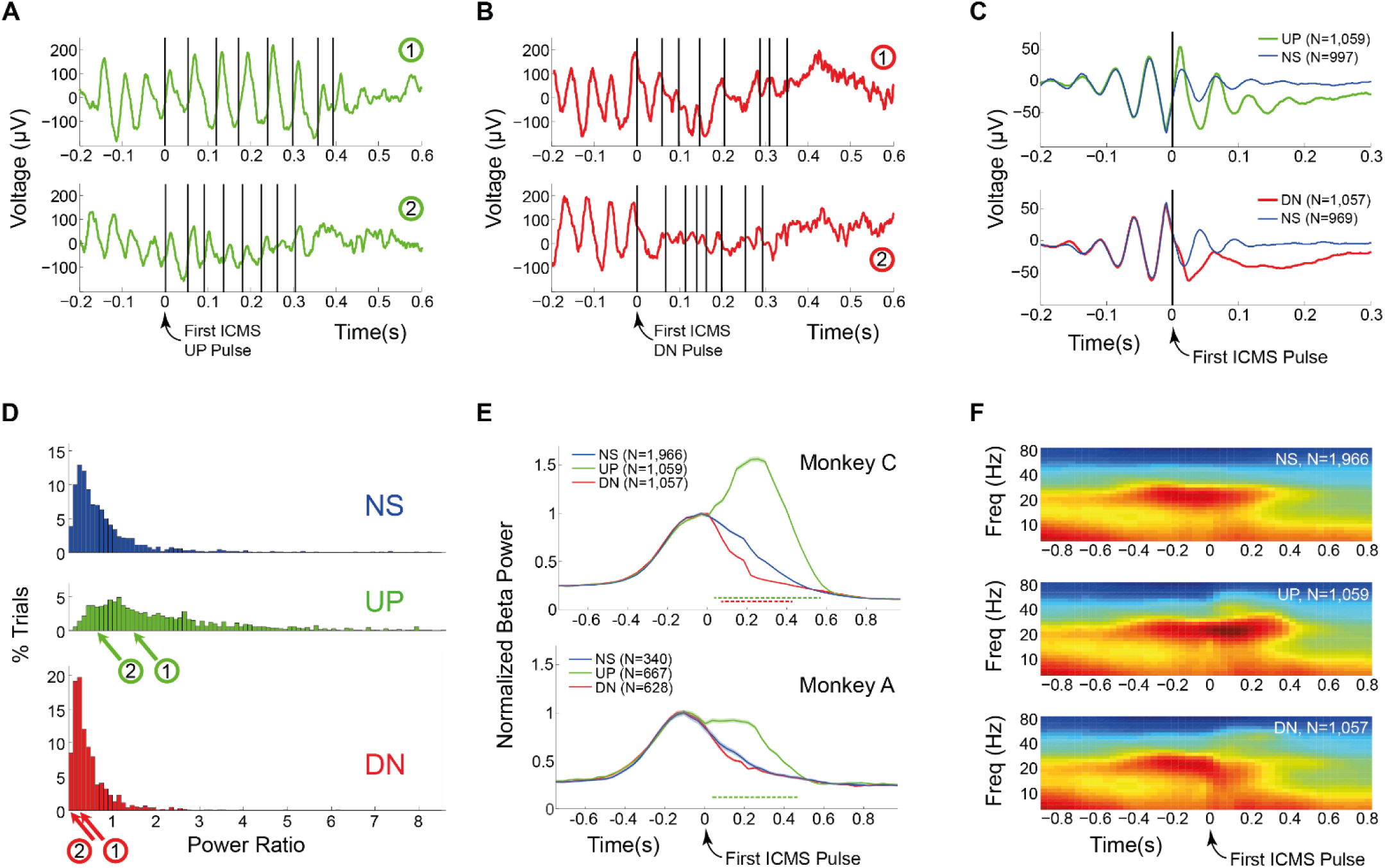
Differential effect of stimulation phase on LFP Beta. (A,B) Single trial examples during UP (green) and DN (red) stimulation. ICMS pulses are marked by vertical black lines. (C) Averaged LFP for UP (top) and DN (bottom) trials. The averaged NS LFP is marked in blue. (D) Histograms of the ratio between Beta-power (20-30Hz) during stimulation and Beta-power at *threshold-reach* for NS, UP and DN trials. Red and green arrows point to the trials shown in (A) and (B). (E) Average Beta-power during stimulation (normalized by the Beta-power at *threshold-reach*) in monkey C (top) and A (bottom). Dashed lines denote the period of significant UP vs. NS (green) and DN vs. NS (red) power differences (p<0.01, Wilcoxon signed-rank test). (F) Spectrograms of the average Beta-power for NS, UP and DN trials. Note the clear increase of Beta by UP stimulation and decrease by DN stimulation as compared to NS. See also Figure S2.

**Figure 6.**
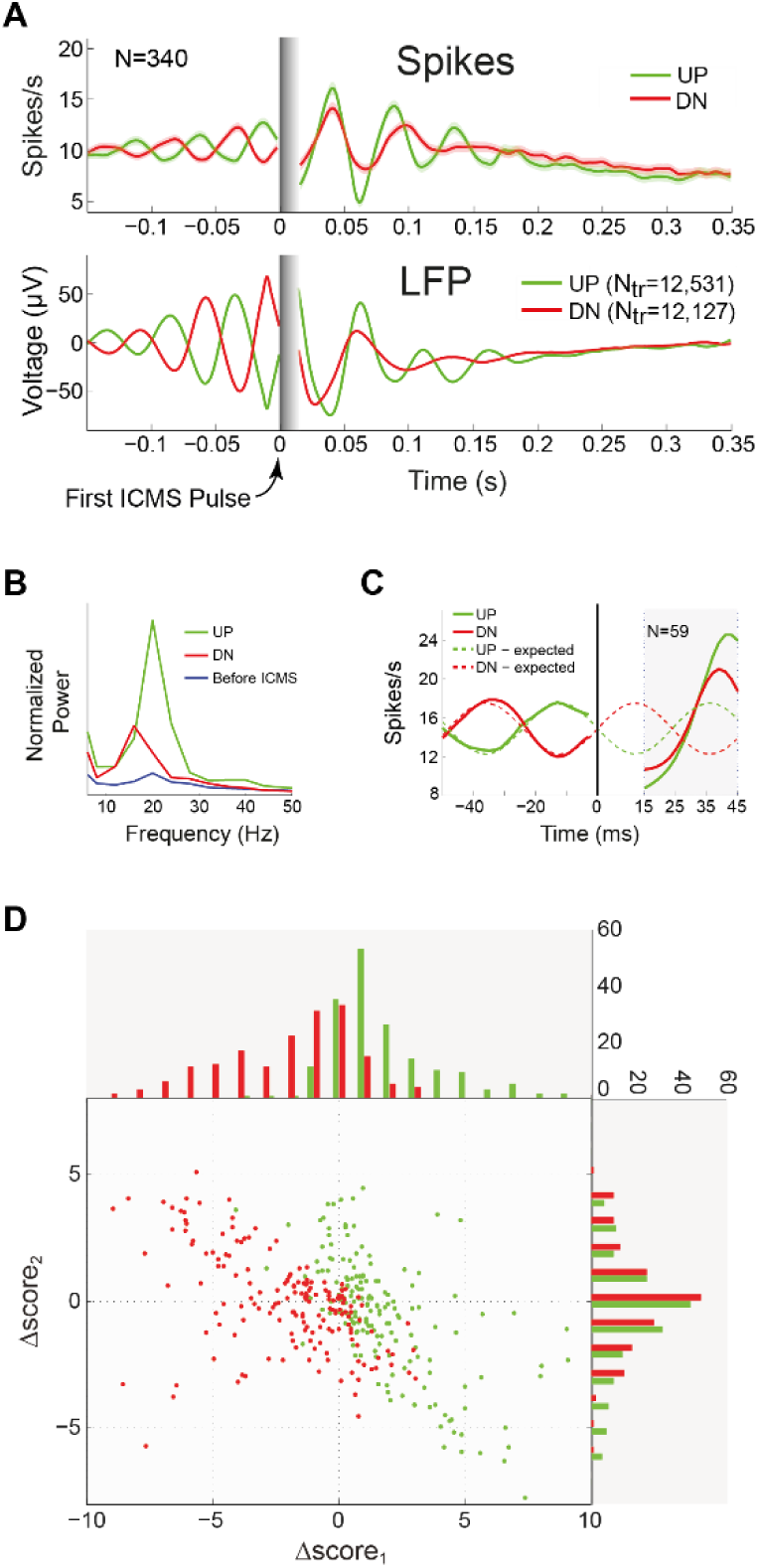
Differential effect of stimulation phase on spiking activity. (A) Mean and SEM of firing rate (top) and LFP signal (bottom) around the first ICMS pulse (time zero) for UP (green) and DN (red) trials. Spike data are based on 340 well-isolated (see Methods) neurons with significant PLI above 0.1. LFP is averaged over the conditioned electrodes. (B) Mean normalized firing rate power-spectrum of the same neurons, computed over 250ms after the first ICMS pulse (UP and DN) and before the ICMS-train. (D) Mean UP and DN firing rate around the first ICMS pulse at time zero, calculated for 59 well-isolated neurons having significant PLI>0.1 and exhibiting prominent pre-stimulus Beta oscillatory activity (see Methods). Solid lines mark the mean UP and DN firing rate, while dashed lines mark the *expected firing pattern*, had there been no stimulation. Shaded gray area marks the 30ms epoch used for the PCA analysis. (D) Comparison of UP Δ*score*_*i*_ (green) vs. DN Δ*score*_*i*_ (red) for neurons recorded within 2mm of the stimulating electrode. *score*_*i*_ is the length of the projection on PC*i* (*i*=1,2) and Δ*score* is the difference between UP or DN score and the projection of the *expected firing pattern*. The distributions of UP and DN Δ*scores* for all neurons are shown in the top (PC1) and right (PC2) histograms. Data are based on 37 sessions. Note that Δ*score*_1_ of UP trials are skewed towards positive values, while the values for DN trials tend to be negative. Notation: N - number of neurons; N_tr_ – number of trials. See also Figures S2, S3 and S4.

Figure 5E depicts the Beta-power as a function of time around the onset of the ICMS-train, normalized by the mean Beta-power at *threshold-reach*. Monkey C showed a high increase in Beta-power for UP trials relative to the NS trials, while in DN trials there was a more moderate, but significant, decrease of Beta. The effects observed in monkey A show a significant, but weaker increase for UP trials with no significant change for DN trials as compared to NS. Note that, for the two monkeys, the oscillatory activity during the UP trials persisted for a longer period after the onset of the ICMS-train, relative to the NS and DN trials.

In addition to the decrease in Beta-power in the DN trials, Figure 5C also suggests a phase shift in the oscillatory wave following the ICMS-train, as the first positive peak of the average DN LFP trace lags the NS trace by almost half the oscillatory cycle. This phase shift can also lead to a drop of the oscillation frequency. To explore this possibility, we computed the mean spectrograms of NS, UP, and DN trials, and found that DN stimulation indeed induce a decrease in Beta-power, accompanied by an increase in lower frequencies power, under 20Hz (Figure 5F).

### Differential effect of stimulation phase on spiking activity

The average spiking activity (Figure 6A, top plot) was affected differentially by the ICMS-train like the LFP (Figure 6A, bottom plot): the oscillatory spiking increased and prolonged following the UP stimuli (green), as compared to the DN stimuli (red). Additionally, DN stimuli resulted in a frequency decrease of the spiking activity (Figure 6B) like the LFP (Figure 5F). The data of Figure 6 include 340 well-isolated neurons with PLI>0.1, recorded by electrodes across the whole array.

We next used principal components analysis to quantify the difference between UP and DN effects on the ongoing oscillatory pattern of spiking activity (see Methods for details). Briefly, we defined a short interval of 30ms after the first (UP or DN) stimulus of each ICMS-train as the *Stimulus evoked pattern* and compared it to the expected patterns in the same interval, in the absence of a stimulus, defined here as *the expected firing pattern*. We then calculated the principal components (PCs) that optimally represent the *expected firing pattern* using a sample of 59 neurons exhibiting prominent pre-stimulus Beta activity in 30ms intervals before the first ICMS pulse. Figure 6C shows the mean firing patterns of these 59 neurons before and after UP (green) and DN (red) stimulation and the mean *expected firing patterns* in solid and dashed lines respectively. Note (in the shaded 30ms post-stimulus interval) that for the UP trials, the *stimulus evoked pattern* (solid green) is in phase with the expected (dashed green) but stronger, and the response to DN stimulation (solid red) is also stronger than expected (dashed red), but phase reversed.

Next, we examined how the *stimulus evoked patterns* of all 340 neurons are represented in the PC-basis. We found that the two first PCs explain over 93% of the variance in these patterns (for details see Figure S3). Interestingly, the representation of UP vs. DN patterns by the two-dimensional PC space is markedly different, as shown by Figure 6D. To create this figure, we defined:

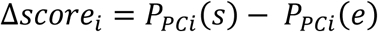

where *s* is the *stimulus evoked pattern* (either UP or DN), *e* is the *expected firing pattern* and *P*_*PCi*_ is the projection score of the *i*_*th*_ principal component (*i*=1,2). Figure 6D shows a scatter plot of Δ*score*_1_ and Δ*score*_2_ for the UP response (green) and DN response (red) for the 166 neurons, which were less than 2mm away from the stimulating electrode (see Supplementary Figure 3 for the rest of the neurons). The top and right histograms in the figure present the distributions of Δ*score*_1_ and Δ*score*_*2*_. The figure shows that 85.5% of the neurons increased their PC1 score following UP stimulation (green dots), implying a strengthening of the *expected firing pattern*, while at the same time DN stimulation caused a decrease in PC1 score in 75.7% of the neurons (red dots), suggesting an interference with the expected oscillatory pattern. Thus, while UP stimulation enhanced the trend of the volitional oscillations, the DN stimulation interfered with this trend.

### The differential effects of stimulation phase decay with distance

Previous studies demonstrated that ICMS at similar currents and higher frequency (200Hz) affect neuronal elements in a radius of 500µm around the stimulation site, and may spread up to 1-2mm depending on the ICMS parameters (Mitz and Wise, 1987; Roe et al., 2015). To evaluate how the stimulation-effects propagate across the array and how they modify LFP signals and single neuron firing patterns, we again used the PCA and applied it to spiking activity and LFP signals recorded by all electrodes of the array (see Methods).

Figure 7 shows spatial maps of Δ*score*_1_ of LFP (top row) and spikes (bottom row) in UP (7A and 7D) vs. DN trials (7B and 7E). The plots show that near the stimulation site there is a significant UP vs. DN differential effect. Namely, the response patterns (of LFP and spikes) follow the general shape of the *expected pattern* during UP stimulation (red zones of the map), while during DN stimulation, the pattern is anti-correlated (blue zones) with the *expected pattern*. Moreover, these plots demonstrate that the effect decays with distance, leaving around half of the array with little or no difference between UP and DN Δ*score*. The differential LFP and spiking activity during UP vs. DN responses and their decay with distance are summarized in Figures 7C and 7F respectively.

**Figure 7.**
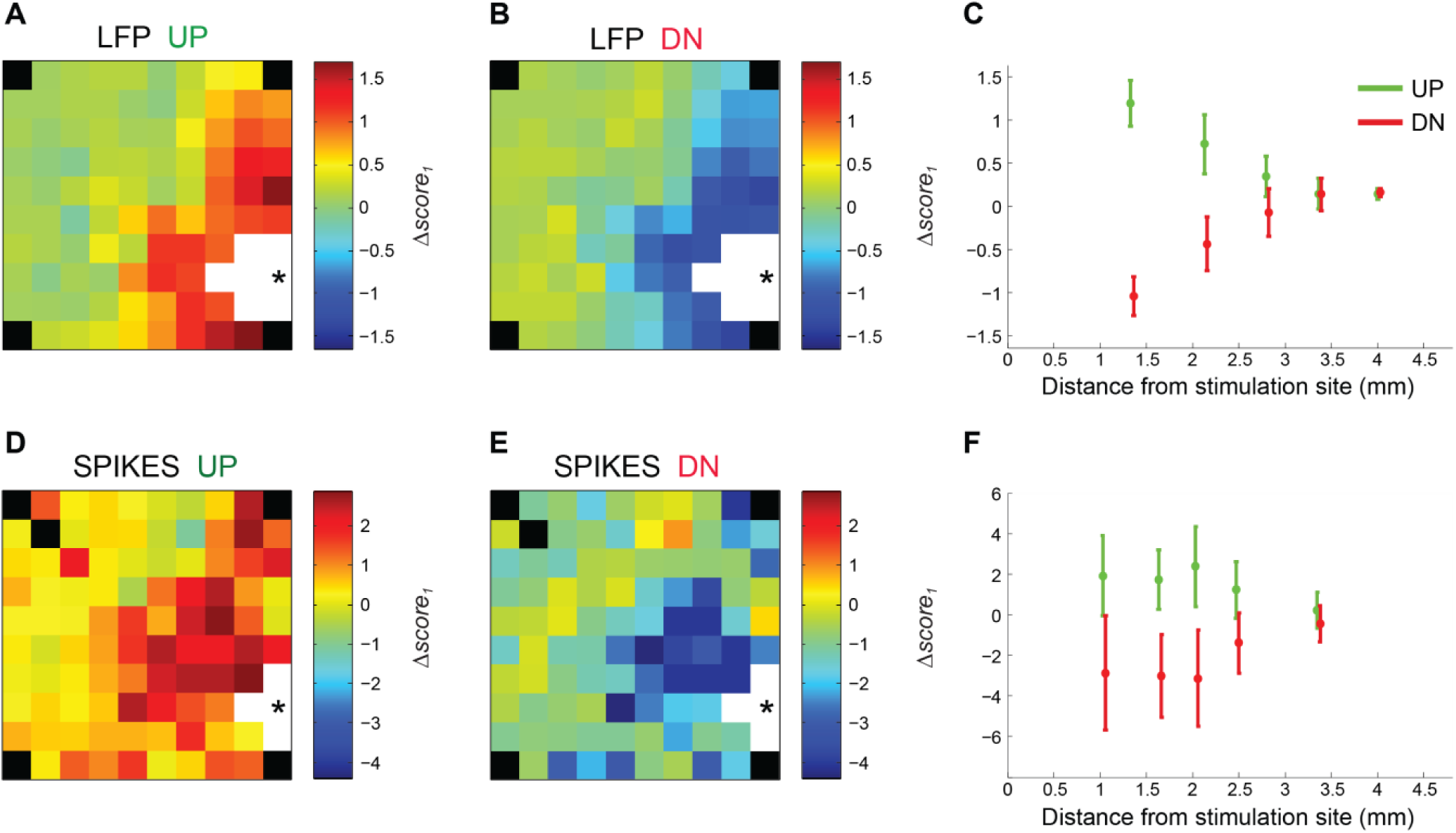
The differential effects of stimulation phase decay with distance. (A,B) Color-coded matrix of LFP Δ*score*_1_ values across the array for UP and DN trials. The stimulation site is marked by an asterisk. White squares denote sites where the recording was occluded by a stimulus-artifact. In black, electrodes that are not connected. (C) Summary of the UP (green) and DN (red) LFP Δ*score*_1_ values vs. distance from the stimulation site. (D-F) same as (A-C) for spiking activity. Data are based on 37 sessions. Note the opposite effects of UP vs. DN stimulation on both LFP and spiking activity: UP stimulus increases the projection on PC1, while DN stimulation decreases it. See also Figures S3 and S4.

### Differential effect of stimulation phase on behavior

So far, we have shown a significant correlation between pre-movement Beta-power and the behavioral indicators (Figure 4). We have also demonstrated that ICMS modulates Beta differentially depending on the phase (UP/DN) of the stimulation (Figures 5, 6 and 7). We next examined the differential effect of the stimulation phase on RT and success rate. Given the positive correlation between Beta-power and RT, we analyzed each of the trial-types (UP/DN/NS) in three regimes of short, medium and long RT, where each regime includes 25% of the trials. Examining RT in these regimes (Figure 8A) revealed that in the trials with long RT, the difference between UP and DN stimulation was statistically significant, exposing a differential effect of ICMS on RT. However, the RTs of the medium and short regimes were not affected by the stimulation. The same phenomenon was observed for the success rate (Figure 8B), namely, only in long-RT trials, the success rate was differentially affected by UP vs. DN stimulation.

**Figure 8.**
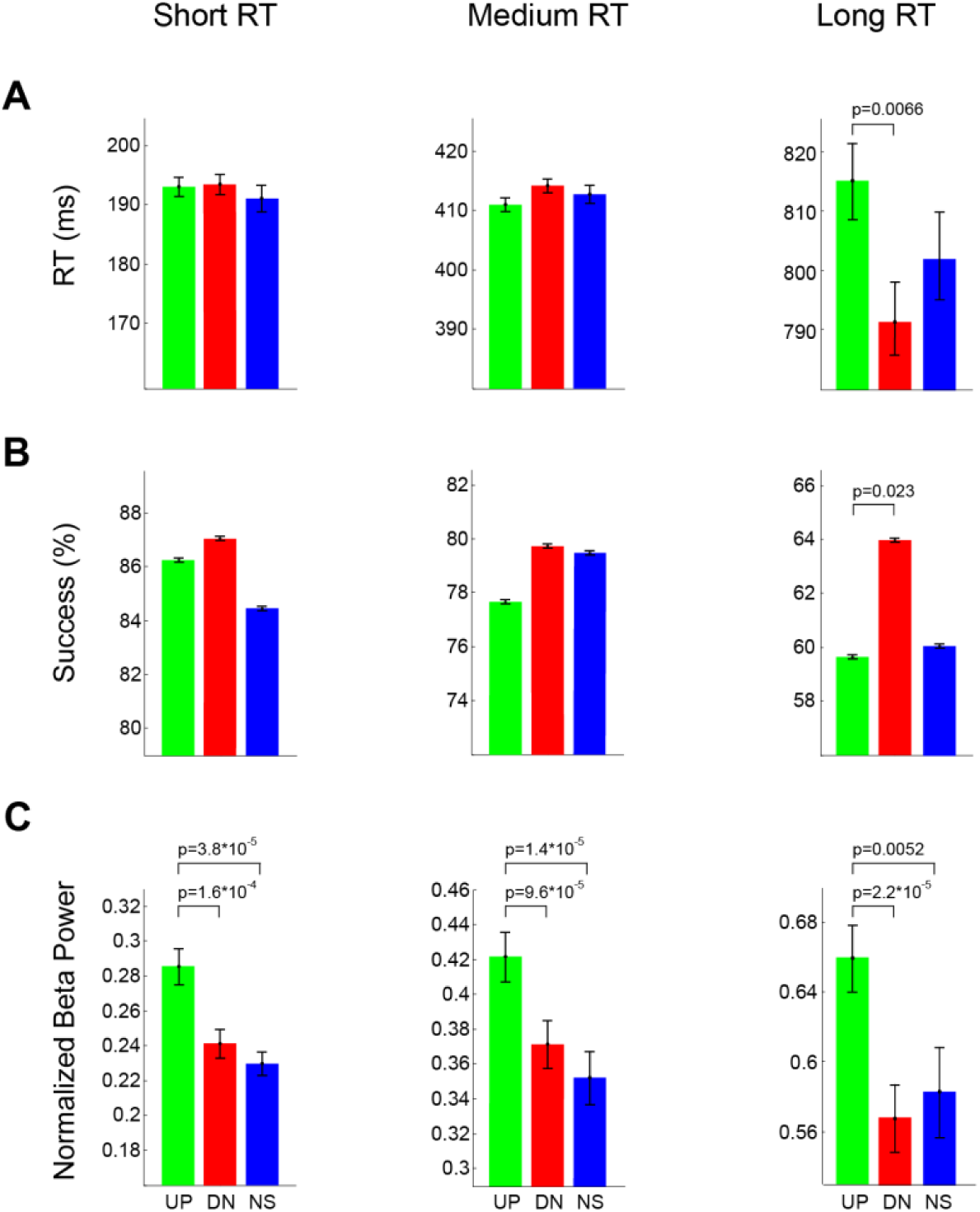
Differential effect of stimulation phase on behavior. (A) RT (mean and SEM) in UP, DN, and NS trials calculated for three regimes of short, medium, and long RT, where each regime includes 25% of the trials. (B,C) Success rate and normalized Beta-power for the same three regimes as in (A). Note that in long RT trials (right column) there is a significant differential effect of UP vs. DN stimulation on RT, success rate and Beta-power. Significance is assessed using Fisher’s test for success rate and Wilcoxon signed-rank test for RT and Beta. Data are based on 30 sessions.

Validating the link between ICMS, Beta and behavior, Figure 8C shows that the Beta-power in these trials was indeed higher in UP trials than in DN trials. Note that the Beta-power was similarly affected by ICMS in all three groups, with a tendency for stronger differential effect in the long RT regime. These results suggest that the effects of ICMS via Beta-power modulation is limited to behaviors that deviate from the normal, expected regime.

## Discussion

This study examines volitional control of LFP and single-unit activity in local cortical circuits, combined with real-time, phase-specific microstimulation as a way to understand the cortical dynamics and its relation to behavior.

Our findings confirm previous work (Khanna and Carmena, 2017; Pogosyan et al., 2009) by showing that increasing Beta disrupts movement initiation and execution. Extending these findings, the study shows that Beta-power is anti-correlated with success rate, suggesting that increased Beta-power also interferes with higher aspects of task performance, including perception and/or decision-making.

We developed a real-time phase-detection algorithm and applied it in our microstimulation procedure. We found that phase-specific ICMS has differential and even opposite effects on volitionally enhanced neuronal oscillations and on spiking activity of single neurons, depending on its precise timing. These effects are site-specific, decaying rapidly with distance from the stimulation site. Lastly, we show that ICMS has phase-specific differential effects on task performance (RT and success rate) in some regimes of behavior.

### Volitional control of Beta Oscillations

Studies of neural conditioning in BMI settings demonstrated in recent years that subjects can volitionally modulate spatiotemporal patterns of activity in local cortical circuits. We confirm previous work (Engelhard et al., 2013; Khanna and Carmena, 2017) by showing that monkeys learn within 2-4 weeks to modulate Beta oscillations. After the initial learning period, their performance reached a new stable state where the Beta oscillations were dramatically higher as compared to the pre-learning state (Figure 2). The high Beta oscillations evolved in each session for several months of recordings in the two monkeys. These learning-induced changes of the local circuit suggest that the BMI conditioning was effective in forming long-term plasticity in the conditioned site.

### Differential effects of UP/DN stimulation on LFP and spikes

The results show that the LFP Beta signal is accompanied by a similar pattern in the firing rate of many neurons, reflecting an oscillatory state of the underlying network. During this state, neurons tend to fire together, expressing synchronized Beta rhythm. Namely, these neurons increase their firing rate at specific phases of the LFP oscillation, while other phases have sparser neuronal firing. Since LFPs are extracellular recordings that reflect mainly averaged subthreshold (e.g., synaptic) activities (Buzsaki et al., 2012), we associate the LFP’s DN and UP phases with “depolarizing” and “hyperpolarizing” phases of the membranes currents respectively.

In our experiment, the main initial effect on LFP in the vicinity of the stimulating electrode was “upward”, towards the hyperpolarization, inhibitory regime, with a net suppression effect of spiking activity. This result is in line with Jackson et al. who demonstrated that antidromic stimulation in the pyramidal tract induces an inhibitory effect that resets the phase of LFP and spiking activity (Jackson et al., 2002). We found the same reset-effect in electrodes that were near the stimulating site (see Figure S4). This stimulus-induced inhibition may explain the differential effects of UP vs. DN stimulation on the oscillatory activity: During UP trials, it increases the hyperpolarizing trend and during DN trials it reduces the depolarizing trend. A possible mechanism to support this explanation is based on the likely assumption that ICMS recruits mainly inhibitory interneurons, due to their higher input resistance and lower threshold. Under this assumption, UP stimulation increases the activity of inhibitory neurons during averaged hyperpolarization of the overall population (80% excitatory neurons), increasing the inhibition and enhancing the oscillatory trend. The opposite happens in DN stimulation, where the inhibitory neurons (which are activated by the stimulus) induce hyperpolarization currents on the population. This effect acts against the overall depolarization of the LFP (DN slope) and reduces the oscillatory trend.

### Effects on behavior

We show that the choice of the stimulation phase had a dramatic differential effect on the neuronal state as portrayed by the LFP and spiking activity in the vicinity of the stimulating site. However, the effects on behavior (RT and success rate) were exposed only in a limited regime: the regime of longest RTs. This apparent discrepancy may be resolved by the hypothesis that the ICMS effects are reflected in behavior only when the circuit loses stability and produces high RT values. In this regime, the stimulation effects on the circuit are indeed reflected in the behavior (Figure 8): UP stimulation enhances the current trend (maintains the phase, increases Beta and increases RT) while DN stimulation disrupts the oscillatory trend (inverses the phase, decreases beta and decreases RT).

The explanation of this hypothesis could be based on the balance between the local activity, which we monitor and affect by the local ICMS, and the activity in wider (remote) circuitry of motor-control, which is predominantly resistant to the local perturbations in our experiment. It is possible that only when the total activity (local + remote) is unstable (producing higher RT and lower success rate) the phase of stimulation (UP/DN) is sufficient to affect behavior differentially. Future experiments are necessary to test this interpretation and extend this line of research, using larger samples of neurons from wider cortical and subcortical brain regions.

### Translational aspects

Following previous studies of volitional neuronal control (biofeedback), we present here a scientific tool, which uses volitional control to harness the inherent brain plasticity, to modulate neuronal state, and affect behavior. Extending these studies, we combine volitional control with fine-tuned electrical stimulation into a hybrid platform that can facilitate restoration of normal activity whenever abnormal patterns are identified.

This hybrid system can be highly beneficial for treating undesired brain oscillations at different frequency bands, which have been evidenced in brain diseases, including Parkinson’s, schizophrenia and others (Brown et al., 2001; Little and Brown, 2014; Raz et al., 2000; Uhlhaas and Singer, 2010). For instance, it could serve as a future closed-loop, deep brain stimulation (DBS) (Little et al., 2013; Priori et al., 2013; Rosin et al., 2011) tool, detecting deviations from normal functioning, inducing volitional control of neuronal activity and injecting phase-locked stimuli, thereby placing the neural activity back on track.

## Acknowledgments

The authors would like to thank A. Shapochnikov and S. Freeman for their technical assistance; M. Doron for programming; B. Engelhard and A. Globerson for their help in planning the experiments; and I. Nelken, M. London, R. Paz and D. Hansel for their helpful comments. This study was supported in part by MAFAT, the Rosetrees Trust, the Gatsby Charitable Foundation, the Adelis Foundation and the Jack H. Skirball Chair & Research Fund in Brain Research. O.P. was supported by Goren-Khazzam scholarship.

## Author contributions

O.P. and E.V. conceived the idea and designed the experiments. Z.I. and E.V. performed the surgeries. O.P. and U.W.R. prepared the setup, trained the animals and collected the data. O.P., H.B. and E.V. analyzed the data. O.P. and E.V. wrote the paper.

## Methods

### Animals and electrode implantation

Two female monkeys (*Macaca fascicularis*, weight 4 kg) were chronically implanted with 10×10 microelectrode-arrays (Blackrock Microsystems) with 400 µm inter-electrode distance in the arm area of M1 of the Left hemisphere (see Figure S1 for array location). Electrodes were coated with iridium oxide and 1.5mm in length. Animal care and surgical procedures were in accordance with the National Institutes of Health Guide for the Care and Use of Laboratory Animals and supervised by the Hebrew University Ethics Committee for the Care and Use of Laboratory Animals.

### Experimental setup

Monkey sat in a behavioral setup, awake and performing a BMI and sensorimotor combined task. A data acquisition and neural stimulation system (AlphaLab SnR, AlphaOmega, Nazareth, Israel) was used for recording LFP and spikes from 96 microelectrodes and applying ICMS to a selected electrode.

The closed loop BMI analyzed the neural data in real time and decided when to reinforce desired pattern of activity. The BMI was embedded in custom-made software for behavioral-control, providing visual and auditory feedback as well as the food reward.

### Behavioral task

Each trial began with a visual cue (orange circle) instructing the monkey to make a small extension of the right (contralateral) palm to express alertness (Figure 1A, stage 0). Next, the monkey was conditioned to enhance the LFP Beta-power in 2-3 adjacent electrodes at a pre-selected location of the array, receiving a visual feedback (green ring of a radius proportional to the Beta-power, around the orange circle) from the BMI algorithm (Figure 1A, stage 1). When a required Beta threshold has been reached, the monkey received a cue (green ring disappeared) and a period of 400-600ms began during which the orange circle turned either lighter or darker (Figure 1A, stage 2). We refer to these colors as “yellow” or “red” respectively. The “yellow” and “red” colors were very similar (RGBs of (255, 136, 0) and (255, 130, 0)), respectively), imposing fixation and attention in order to discriminate between the colors. At this epoch an ICMS burst of around 300ms was delivered in 80% of the trials through the selected electrode. During each session, the threshold could be manually adjusted so as to maintain a success rate of around 75-80%.

Upon receiving a visual Go-signal (circle turned green), the monkey had to decide if the circle is yellow or red and report by pressing one of two keys with the contralateral hand (Figure 1A, stage 3). Food reward and auditory positive-feedback were delivered based on the monkey’s report and the Beta-power after threshold had been reached. If the power remained above a threshold ß1, correct responses were always rewarded, and if the power was below ß1 and above ß2, correct trials were randomly rewarded with probability proportional to the Beta-power. Wrong key press trials or trials with beta below ß2 were not rewarded and an auditory negative-feedback was delivered. The monkey’s hand movements were monitored via an accelerometer attached to the middle finger. This was used both for measuring reaction time and for automatically failing the trial when a movement occurred during the conditioning or stimulation epochs of the trial (Figure 1A, stages 1 and 2).

### Session structure and data acquisition

Each recording session was composed of 12-24 alternating blocks of behavioral trials (~150±34) and control trials (46±10). In the latter, no behavior was required (Figure S2) and rewards were solely dependent on the Beta-power. LFP signals were sampled at ~1.4 KHz and filtered using a 0.1–250 Hz band-pass filter. Spiking activity was sampled at ~22.3 KHz, filtered using a 0.25-8 KHz band-pass filter and sorted online by the SNR recording system.

### Conditioning algorithm

The conditioning data were based on the LFP signal of 2-3 adjacent electrodes in a selected location where Beta was evidently observed before we started the conditioning sessions. The location of the conditioning electrodes was slightly adjusted during the first 26 and 19 sessions of the learning period for monkeys C and A, respectively.

The BMI algorithm calculated for each conditioning electrode the mean power in a band between 20-30Hz (“Beta”), using the average of the discrete Fourier transform (DFT) coefficients in that frequency band. This value was computed every 50ms on a 333ms window. The mean power of the 2-3 electrodes was used for displaying the visual feedback (green ring), and for detecting Beta-threshold crossings (like *threshold-reach* in Figure 1).

To achieve a smoother, more natural motion of the green ring, a smoothing window was applied over every 3 consecutive Beta-calculations. In addition, a maximal step size was set to prevent jitter of the display. To impose volitional control of Beta without increase in the lower frequencies, an upper limit was set over a band between 8-16Hz. Once this limit was reached, the movement of the ring temporarily froze.

### Electrical stimulation and real-time phase detection

ICMS-trains were applied through one electrode, selected to be in close proximity to the conditioning electrodes. In each trial, stimulations began after reaching the desired threshold (Figure 1A, end of stage 1). The timing of each pulse was determined as follows: the LFP data from a single conditioning electrode were filtered by a Savitzky-Golay smoothing and differentiation filter (Savitzky and Golay, 1964), and a zero-crossing detector was applied to the derivative to reveal the signal’s peaks. If the detected peaks matched the required Beta frequency (20-30Hz), ICMS was applied on either the rising phase (UP trials) or falling phase (DN trials) of the oscillation, aiming at about one-third of the rise or fall, respectively. The peak detection algorithm continued after each pulse, applying the next pulse no less than 25ms and no more than 120ms after the previous one. The time between the first and the last pulses was limited to 400ms, hence each burst consisted of 4 to 8 pulses (5 or more pulses in 99.95% of the trials). The pulses were cathodic-phase-first with a phase of 100µs and intensity of 80µA (with some controls at 40, 60 and 120µA). Each ICMS-pulse induced an artifact of around 12msec, during which we could not detect spikes activity or LFP. For the LFP we estimated the voltage by spline interpolation.

During some sessions, the control trials consisted of a single ICMS pulse. This pulse was applied at a random phase of the beta oscillation (Figure S2).

### Data Analysis

All post-processing was performed in MATLAB (MathWorks). The power spectrum was calculated using the DFT coefficients in segments of 350ms (Figure 2). For Beta-power calculation, only those coefficients between 20 and 30Hz were used (Figures 1,2,4,5, and 8). For Figures 5E and 5F, we used somewhat shorter segments of 250ms with an overlap of 215ms to illustrate the evolvement of Beta as a function of time.

Single-units were sorted online by AlphaLab SnR. For some analyses, all neurons were used, while for others, only neurons with isolation scores above 0.74, and a signal to noise ratio (SNR) above 7 were selected (*well-isolated neurons*). Isolation score was determined by a modified version of an isolation quality assessment procedure (Joshua et al., 2007).

### Behavioral indicators

Success rate was defined as the percent of trials, which ended in a correct key press. Note that this is different from the rewarded trials as the monkey did not receive reward for trials in which the Beta-power was too low after *threshold-reach*. Wrong key press was the predominant cause of trial failure (99%). The reaction time (RT, from Go-signal to movement initiation) was determined based on the signal of the accelerometer. In the behavioral analysis, only trials in which the monkeys pressed the left key were considered. To calculate the movement time (MT, from movement initiation to key press), RT was subtracted from the total time, from Go-signal to key press. For the computation of behavioral indicators as a function of Beta (Figure 4), the Beta-power of each trial was normalized by the mean daily Beta-power calculated over the 350ms prior to *threshold-reach*. The normalized power values were then sorted and filtered by a moving average procedure using a Gaussian window (length: 25% of the number of power values, σ: 8%, window-overlap: 96%) producing a Beta-power vector. The mean and SEM of the behavioral time indicators (RT, MT, RT+MT) were then computed for every element in this vector, based on the data of the corresponding trials. The mean success rate was computed in the same way as the behavioral time indicators. SEM of the success rate was calculated by bootstrapping (1000 random sampling of the original set size, with replacements).

### Phase locking index (PLI) and preferred phase

We used a phase locking index (Lachaux et al., 1999) to quantify the level of synchrony between the firing of each neuron and the Beta. To do that, we used the LFP of each electrode during the 200ms preceding *threshold-reach* (which presumably consist of high Beta-power) and selected the third of the trials with the highest Beta-power. We then filtered the LFP signal in the Beta range (20-30Hz) and applied the Hilbert transform to the filtered signal. The phase of the Hilbert transform was used as the instantaneous phase of the LFP signal at the Beta frequency. Next, each spike was assigned a phase corresponding to the instantaneous phase of the LFP recorded from the same electrode during the time of the spike. Finally, we performed vector-averaging of each neuron’s phases to get its mean phase vector. The PLI is the size of this vector and the preferred phase of the neuron is the vector’s phase. Mathematically the PLI of a specific neuron can be computed as follows:

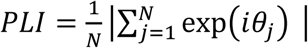

Where *θ*_*j*_ is the instantaneous phase of the j^th^ spike and N is the number of spikes of the neuron.

In our analysis, we’ve only used those neurons having statistically significant PLIs. For neuron X having *PLI*_*x*_ and N spikes we randomly generated 1000 vectors of N phases and calculated the PLI of each vector. A neuron was considered to have a statistically significant PLI if the probability of a PLI above *PLI*_*x*_ was less than 0.01.

### PSTH

PSTHs of spiking activity were calculated for all neurons, around the time of the first ICMS pulse, with 2ms bin-size. The PSTHs were then passed through a Gaussian filter of length 7 (14ms) and σ =2 and averaged over all relevant neurons (Figures 6, S2, S3 and S4).

### PCA

To study the differential response to UP and DN stimuli of spiking activity and LFP, we explored the 30ms intervals, starting 15ms after the first ICMS pulse, using PCA. For the spiking activity, we analyzed 340 well-isolated neurons with PLI>0.1. We chose 59 neurons exhibiting prominent pre-stimulus Beta activity, namely, at least 60% of their PSTH variance, during 60ms before stimulation, could be explained by a sine wave of 18 to 30Hz. We used the PSTHs of these neurons to calculate the PCA of the *expected firing pattern*, under the assumption that without stimulation the oscillatory activity persists for another 50ms. We then projected the PSTHs of all 340 neurons after the first ICMS pulse on the *expected firing pattern* PCs to get the UP and DN *score* in PC-space, subtracted the *expected pattern* of each neuron and compared the results of UP vs. DN Δ*score* (Figure 6D and S3). For the spatial maps (Figures 7D and 7E) we used a larger pool of neurons (N=2885) without restricting the isolation score or the PLI, thus enabling better statistics for all electrodes across the array. For each electrode, the final Δ*score* was the median of the Δ*scores* of all neurons recorded from that electrode. In the summary graph (Figure 7F) all neurons were divided into 5 equal-size groups based on their distance from the stimulating electrode.

For the PCA of the LFP, *expected pattern* and Δ*scores* for UP and DN were similarly calculated, based on the LFP activity in NS trials (Figures 7A-C).

