## Supplementary information for "Phase-specific microstimulation in brain machine interface setting differentially modulates beta oscillations and affects behavior"

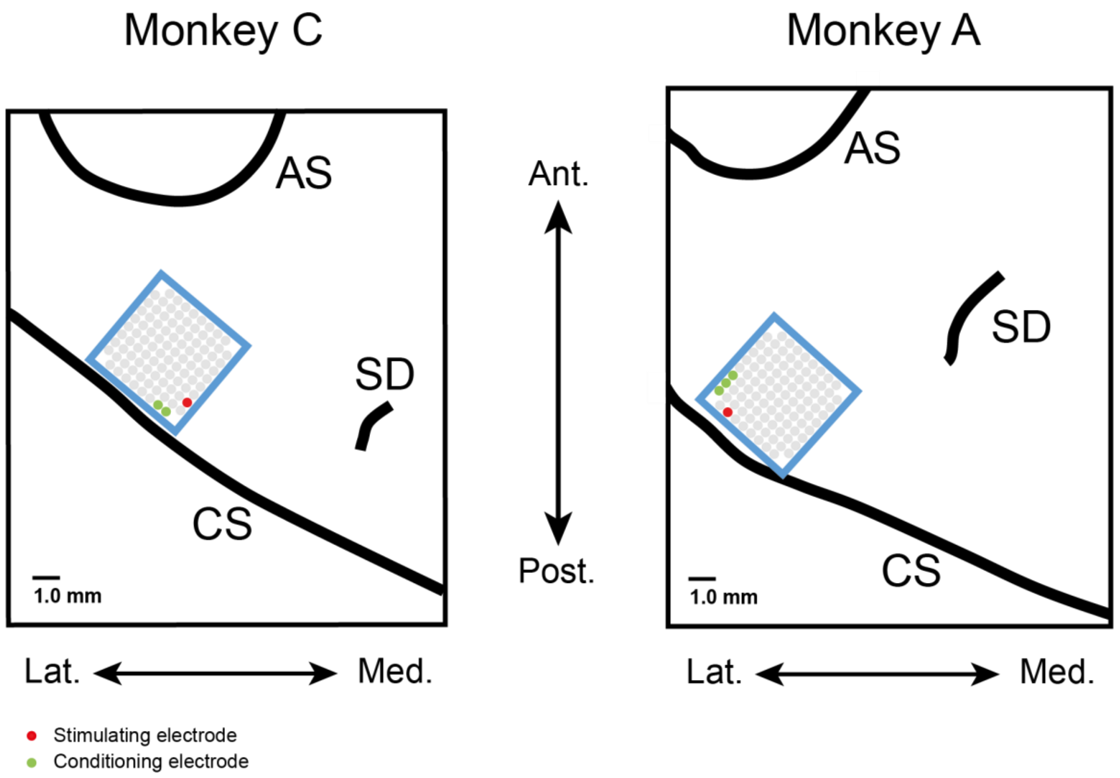


**Figure S1. The Arrays location**

Arrays location in motor cortex for monkey C and monkey A. Red and green dots show the stimulation and conditioning sites during the sessions associated with Figure 3 through Figure 8. Notation: CS - central sulcus; AS - arcuate sulcus; SD- superior dimple.

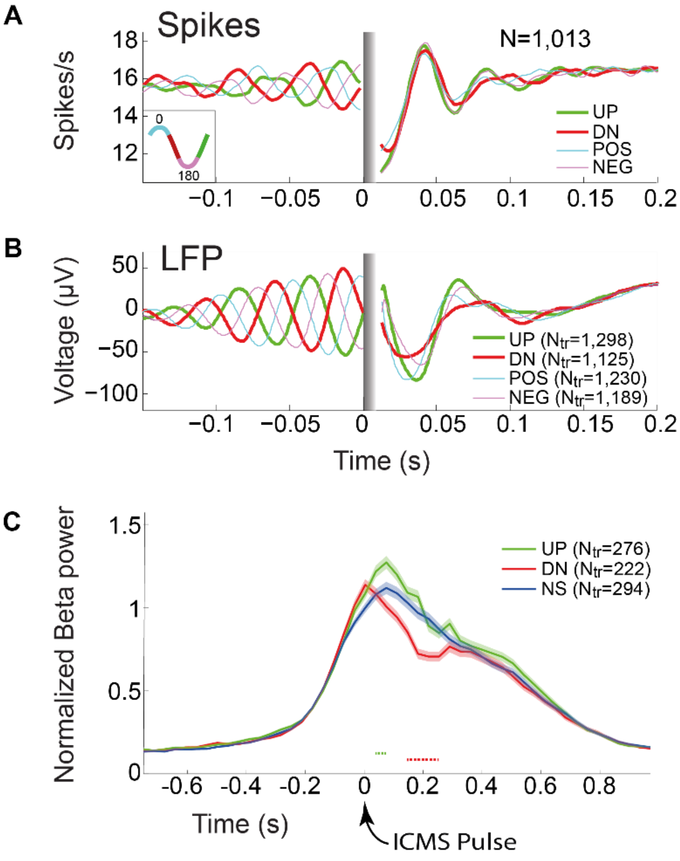


**Figure S2. Single pulse ICMS: UP vs. DN Differential effects on LFP and spiking activity (related to Figures 5 and 6)**

To further examine the choice of UP and DN as the two explored phases in our study, we analyzed the effects of a single-pulse stimulation, at different phases of the oscillations. Here, the stimulus was applied at a random phase, after threshold-reach. We divided all ICMS phases into four groups; two around the positive and negative slopes of the oscillations (resembling UP and DN stimulation phases), and two, around the positive and negative peaks. The four phase-regions are shown by a cycle plot in the inset at the bottom-left of (A).

(A) and (B) present the mean traces of single-unit firing rates and LFPs (recorded by the conditioning electrodes) respectively. Time zero is the time of the ICMS pulse. Note that UP and DN produce the two extreme effects on LFP oscillations as compared to positive

(POS) and negative (NEG) peaks. UP induces the strongest increase of the oscillations, while DN is most effective in reducing the oscillation (the chosen phases for the detailed study).

(C) Average Beta-power around a single ICMS pulse (time zero), showing a relatively short and small ICMS effect. Dashed lines denote the period of significant UP vs. NS (green) and DN vs. NS (red) differences ($p<0.01$, Wilcoxon signed-rank test). The small effect suggests that numerous consecutive stimuli are required to elicit a more substantial increase to the Beta-power, as shown in Figure 5. Notation: N - number of neurons; $N_{tr}$ – number of trials.

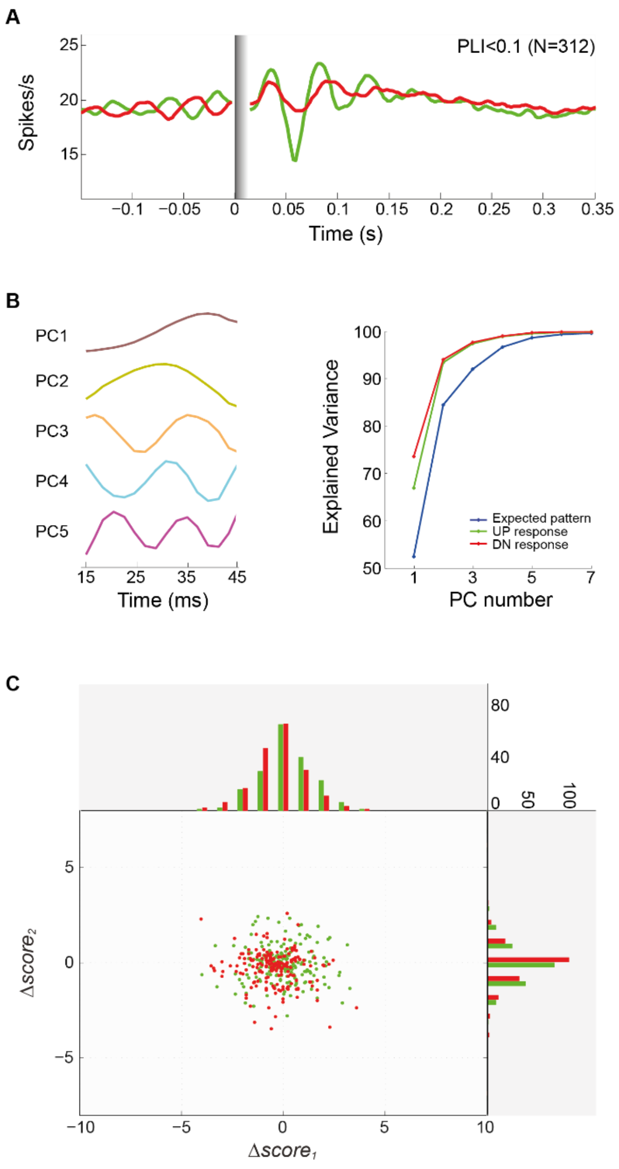


**Figure S3. Differential effect of stimulation phase on spiking activity (related to Figures 6 and 7)**

(A) Mean firing rate of neurons with PLI<0.1. Comparing it to neurons with PLI>0.1 (Figure 6A, top) shows that UP ICMS facilitate oscillations even in neurons with low PLIs, while DN stimulation is less effective in inducing oscillations in these neurons.

(B) PCA analysis. Left: Top 5 PCs of the *expected firing pattern* 15-45ms after the first ICMS pulse. Right: Accumulating explained variance by the top 7 PCs, for the *expected firing patterns* (blue) of the 340 neurons of Figure 6A. Their UP and DN *stimulus evoked* *patterns* are shown in green and red respectively. Note that the first two PCs explain over 90% of the variance.

(C) Comparison of UP $\Delta score_i$ (green) vs. DN $\Delta score_i$ (red) for distant neurons (more than 2mm away from the stimulation site) as shown in Figure 6D for neurons in the vicinity of the stimulation site. The analysis here shows that the distant neurons exhibit overlapping $\Delta score$ distributions, indicating that unlike the nearby neurons (Figure 6D), the distant ones show no differential responses to UP vs. DN stimulations. The combined result is in accordance with the analysis in Figure 7, demonstrating that differential effects of UP/DN ICMS decay with distance from the stimulating electrode.

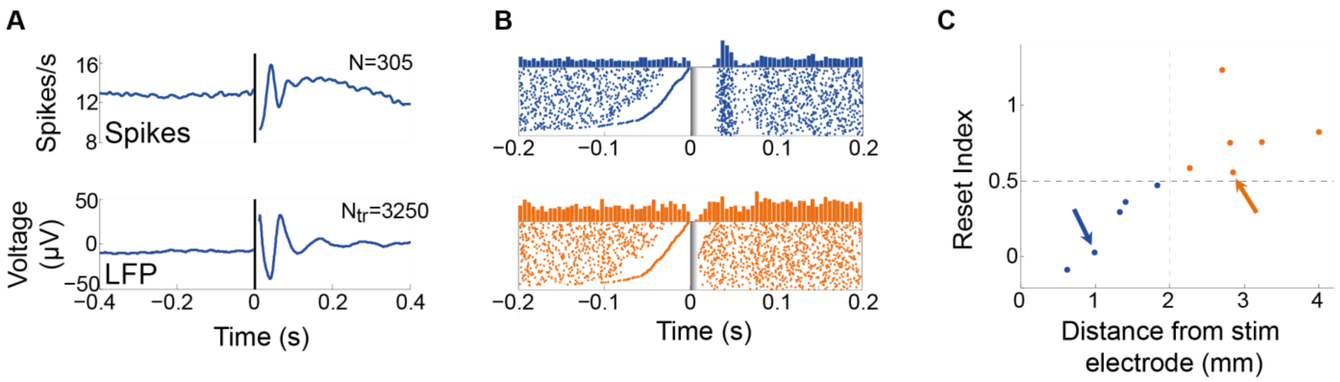


**Figure S4. Distance-dependent phase resetting (related to Figures 6 and 7)**

(A) Average single-units firing-rate (top) and LFP (bottom) around the time of a single ICMS pulse (time 0). Since the stimuli were applied at a random oscillation-phase, the pre-stimulus oscillations were averaged out, and the reset-effect of the stimulus on the oscillation becomes evident, for both firing-rate and LFP activity. As discussed in the paper, the data suggests that each stimulus induces some hyperpolarization, regardless of the phase at which it is applied. During UP trials, this hyperpolarization follows the oscillatory trend, hence the oscillation is enhanced, whereas in DN trials, the hyperpolarization opposes this trend, disrupting the oscillation (see Figure 6). Thus, the reset phenomenon could be a manifestation of the hyperpolarization effect by the stimulating electrode. Notation: N - number of neurons; $N_{tr}$ – number of trials.

(B) Rasters and PSTHs of a near-by neuron (top) and a distant neuron (bottom) relative to the stimulating electrode. The trials in the rasters are sorted according to the last spike before the ICMS pulse (time 0). Note that the silent periods preceding the “last spikes” reflect the Beta firing pattern of both near and distant neurons.

(C) *Reset index* (0 indicates “ideal-reset” and 1 is “no-reset”, see details below) as a function of the distance from the stimulating electrode. Each point shows the average for units at a given distance. The coloring is based on k-means clustering of reset-indices (k=2). Arrows point to the groups containing the two units presented in (B). Data is based on 11 sessions.

Note the distance-dependent reset effect of the stimulation: Firing of nearby neurons tend to undergo phase-reset following the ICMS, whereas distant neurons tend to maintain their phase. Similar decay with distance was presented in Figure 7 for the differential UP vs. DN effect.

*Reset index* calculation: For each trial, we measured the time between the ICMS pulse and two spikes: the last spike before it (T1) and the first spike after it (T2). We omitted all trails with T2 > 60ms (no spikes in the first Beta cycle after the ICMS). We calculated the linear regression of T2 vs. T1 and gradually omitted distant outliers until achieving a regression with $R^2$ of at least 0.4. The *reset index* was defined as the slope of the linear regression of the remaining trials. In Supplementary Figure 4C neurons with *reset index* based on less than 15 trials or under 20% of the original number of trials or PLI<0.05 were omitted. In addition, only points with a minimum of 5 neurons are included in the graph.
